# Tabula-rasa exploration decreases during youth and is linked to ADHD symptoms

**DOI:** 10.1101/2020.06.11.146019

**Authors:** M Dubois, A Bowler, ME Moses-Payne, J Habicht, N Steinbeis, TU Hauser

## Abstract

During childhood and adolescence, exploring the unknown is important to build a better model of the world. This means that youths have to regularly solve the exploration-exploitation trade-off, a dilemma in which adults are known to deploy a mixture of computationally light and heavy exploration strategies. In this developmental study, we investigated how youths (aged 8 to 17) performed an exploration task that allows us to dissociate these different exploration strategies. Using computational modelling, we demonstrate that tabula-rasa exploration, a computationally light exploration heuristic, is used to a higher degree in children and younger adolescents compared to older adolescents. Additionally, we show that this tabula-rasa exploration is more extensively used by youths with high attention-deficit/hyperactivity disorder (ADHD) traits. In the light of ongoing brain development, our findings show that children and younger adolescents use computationally less burdensome strategies, but that an excessive use thereof might be a risk for mental health conditions.

## Introduction

Despite limited knowledge and cognitive capacity, children are known to be very good learners (Gopnik, Griffiths, and Lucas 2015; Kidd and Hayden 2015). Understanding how they achieve such rapid learning is the holy grail of artificial intelligence (Turing 1950), and could help identifying developmental disorders that suffer from learning impairments (e.g. ADHD, dyslexia or dyscalculia; Kaufmann and Aster 2012; Luman, Tripp, and Scheres 2010; Snowling 2014) Central to efficient learning is the exploration-exploitation dilemma (Kidd and Hayden 2015; Sutton and Barto 1998), where an agent needs to arbitrate between exploring less known options to make better informed (and possibly more rewarding) decisions in the future and exploiting the current best known option for higher immediate reward.

Recent studies (Gershman 2018; Schulz and Gershman 2019) have shown that adults use at least two strategies for exploration: One which is based on the injection of stochasticity (captured by Thompson sampling; Thompson, 1933) and one which is directed towards information gain (captured by Upper Confidence Bound, UCB; Auer, 2003). However, both those methods require individuals to keep track of both the expected values and the uncertainty associated to each bandit (Dubois et al. 2020; Gershman 2018). This makes them expensive in terms of resources and thus not always usable (Dezza, Cleeremans, and Alexander 2019; Dubois et al. 2020). This limitation might be particularly pertinent in children and adolescents given their reduced cognitive and neural capacity, which is at least partially due to the delayed maturation of higher cognitive areas, such as the prefrontal cortex (PFC; Casey, Tottenham, Liston, & Durston, 2005; Hartley & Somerville, 2015; Ziegler et al., 2019). We have recently shown (Dubois et al. 2020) that even adult humans deploy two simple heuristic exploration strategies in addition to the computationally expensive strategies: tabula-rasa and novelty exploration. Tabula-rasa exploration (captured by *ϵ* -greedy; Sutton & Barto, 1998) is the cheapest way to explore whereby the same probability is assigned to all options, essentially ignoring all prior information. Novelty exploration is another heuristic whereby an option not encountered previously is chosen. This captures the intrinsic value of choosing something new by adding a novelty bonus (Krebs et al. 2009) to previously unseen choice options.

Exploration is thought to be high in children and to diminish as they grow older (Gopnik 2020; Gopnik, Griffiths, and Lucas 2015), which stands in stark contrast to the limited resources that developing youths have for sophisticated problem solving. A solution for this paradox could be that they use (phylogenetically old) heuristic strategies, which may not be optimal but efficient under given constraints. Experimentally, however, evidence for this hypothesis is limited. Previous studies have found differences in computationally complex strategies such as a change in the valuation of uncertainty in UCB exploration (Schulz et al. 2019) and a change in functional exploration (Somerville et al. 2017) between children and adults, but none have considered the utilisation of simpler and more straightforward heuristics, and how they develop before adulthood. We hypothesize that tabula-rasa exploration, or in other words computationally cheap exploration without integrating prior knowledge, plays a particularly crucial role at a young age.

Understanding the developmental trajectories of exploration strategies may also be relevant for understanding developmental psychiatric disorders. We have previously shown that excessive exploration is a mechanism underlying attention-deficit/hyperactivity disorder (ADHD; Hauser et al. 2014, 2016). However, previous work was unable to pinpoint the specific exploration strategy that goes awry in ADHD. Since then, pharmacological manipulations have demonstrated that tabula-rasa exploration is modulated by noradrenaline (Dubois et al. 2020), which is one of the neurotransmitters believed to contribute to ADHD (Arnsten and Pliszka 2011; Berridge and Devilbiss 2011; Del Campo et al. 2011; Frank et al. 2007; Hauser et al. 2016). We thus hypothesize that tabula-rasa exploration may be excessive in those with high levels of ADHD symptoms.

To test our hypotheses, we used a child-friendly apple gathering task which allows us to tease apart the contributions of complex exploration strategies and exploration heuristics. Using behavioural markers and computational modelling, we show that both younger age groups as well as those scoring higher on ADHD symptoms display an increased utilisation of tabula-rasa exploration.

## Methods

### Subjects

We recruited 108 subjects from schools in Greater London. 11 subjects were excluded from the analysis: 10 due to incomplete data collection (Cogent is not Windows 10 friendly) and 1 due to a medical condition. The final sample consisted of 26 children (16 female; age: M=9.32 years, SD=.27, range=8.89-9.71), 38 young adolescents (21 female; age: M=13.13 years, SD=.30, range=12.69-13.64), and 33 old adolescents (19 female; age: M=17.18 years, SD=.29, range=16.71–17.45). Age groups did not differ in gender nor intellectual abilities (cf. Table S1 in Supplementary Materials). Each subject was given a gift voucher of £7.

As a measure of ADHD symptoms, we used the Self-Report Conners 3 ADHD Index (Conners 3AI-SR, adjusted for age and gender, Conners 2008). The Conners 3AI-SR is an index which contains the 10-items from the full-length Conners 3 questionnaire which best differentiate youths with ADHD from youths in the general population (Conners 2008). All age groups had similar ADHD scores (cf. Table S1 in Supplementary Materials). All subjects provided written informed consent and everyone below age 16 provided written permission from a parent or legal guardian. The study was approved by the UCL research ethics committee.

### Task

We used a multi-round three-armed bandit task (Fig. 1a; bandits depicted as apple trees) in which subjects were asked to maximise an outcome (i.e. amount of juice proportional to the sum of apple sizes) by choosing between bandits carrying varying levels of information. The task design allows for the assessment of the contributions of tabula-rasa and novelty exploration heuristics in addition to the complex Thompson sampling and UCB exploration (Dubois et al. 2020). Taking advantage of the fact that both heuristics make specific predictions about choice patterns, in the “Maggie’s Farm” task, rewards (i.e. apple size) and prior information (i.e. initial samples) about bandits were carefully crafted to capture meaningful variation (Dubois et al. 2020). Bandits carry either a lot, some or no prior information (i.e. 3, 1 or 0 initials samples). Novelty exploration assigns a ‘novelty bonus’ only to bandits for which subjects have no prior information, predicting to choose it disproportionally more often. In contrast, UCB assigns a bonus to each bandit proportionally to how informative they are. Bandits had either a high, medium or low reward sampling mean (with fixed sampling variance). Tabula-rasa exploration predicts that all prior information is discarded entirely and that there is equal probability attached to all choice options, predicting to occasionally choose bandits known to be substantially worse than others. Such known poor value options are not favoured by any other exploration strategies, even by what is traditionally known as ‘random exploration’ (Gershman 2018; Wilson et al. 2014), which is captured by our UCB model (for discussion cf. Dubois et al. 2020).

**Fig. 1.**
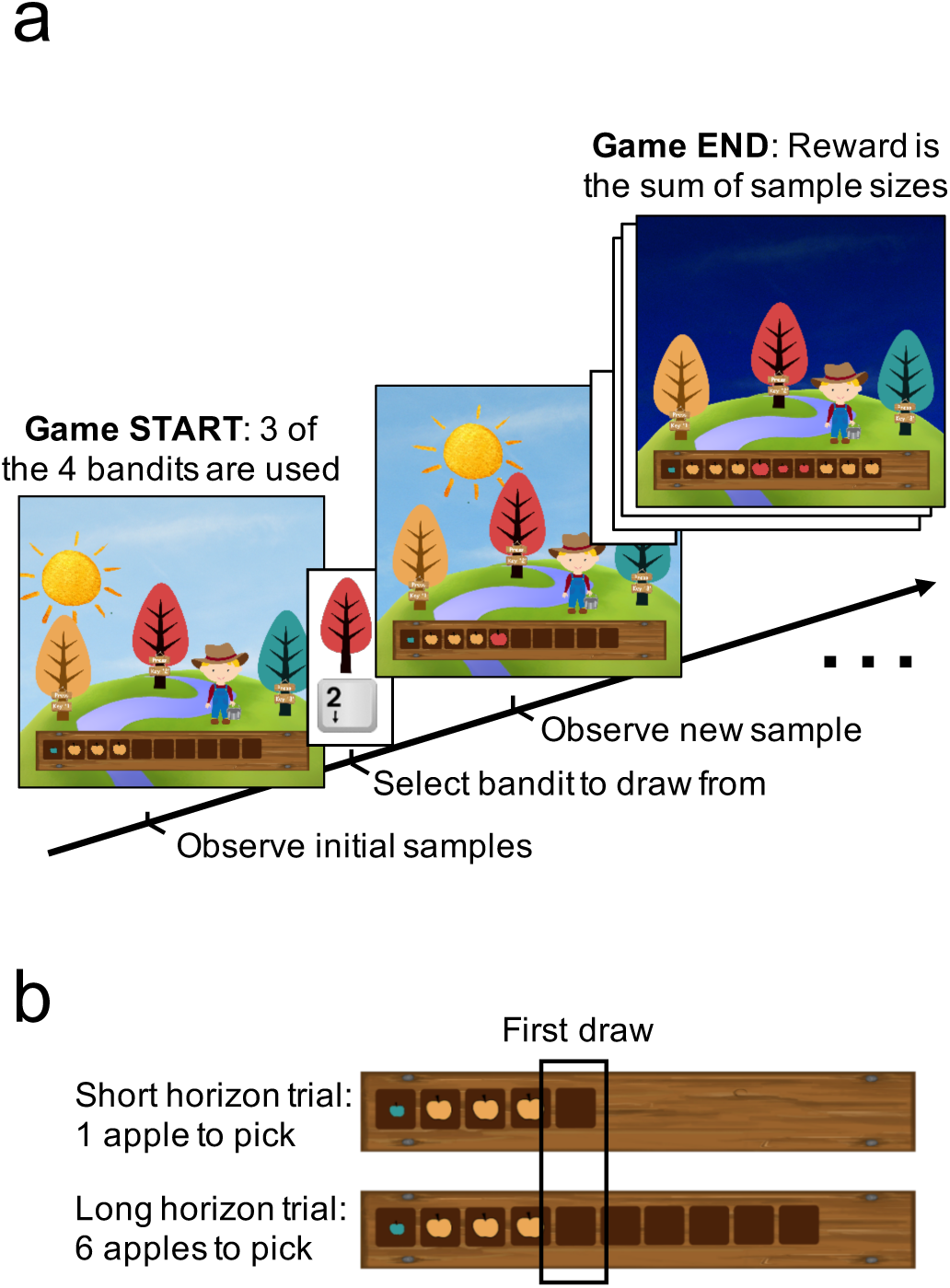
Study design. In the Maggie’s farm task (Dubois et al. 2020), subjects had to choose from three bandits (depicted as trees) to maximise their overall reward. The rewards (apple size) of each bandit followed a normal distribution with a fixed sampling variance. (a) At the beginning of each trial, subjects were provided with some initial samples (number varied depending on the bandits present on that trial) on the wooden crate at the bottom of the screen and had to select which bandit they want to sample from next. (b) Depending the condition, they could either perform one draw (short horizon) or six draws (long horizon). The empty spaces on the wooden crate (and the sun position) indicated how many draws they had left.

In the analysis, we divided these bandits into three different types. The ‘high-value’ bandit was the bandit with the highest sampling mean on a given trial and carried either a lot or some prior information (i.e. 3 or 1 initial samples). The ‘low-value bandit’ carried some prior information (i.e. 1 initial sample) from a substantially lower generative mean, thus appealing to a tabula-rasa exploration strategy alone. The ‘novel bandit’ carried no prior information (i.e. 0 initial sample) and therefore appealed to a novelty exploration strategy.

The degree to which subjects could interact with the same bandits (similar to previous exploration tasks; Wilson et al. 2014; Fig. 1b) was manipulated to modulate exploration. On each trial, subjects could make in total either one draw, encouraging exploitation (short horizon condition) or six draws encouraging exploration (long horizon condition). The key point of this manipulation is that in the long horizon subjects have more opportunity to benefit from the information gain that was obtained through exploration, whereas in the short horizon, the gained information cannot be put to use and is therefore futile. Subjects performed a training session to make sure they understood the instructions and the general concept before playing a total of 96 trials (48 in each horizon condition) during the task.

### Statistical analyses

We compared behavioural measures and model parameters using repeated-measures ANOVAs with the age group as between-subject factor (children, young adolescents, old adolescents) and the decision horizon as within-subject factor horizon (long: 6 choices, short: 1 choice). We report effect sizes using partial eta squared (η^2^) for ANOVAs and Cohen’s d (d) for t-tests. To assess correlations between ADHD and exploration strategies, we z-scored all measures and exploration strategy parameters were averaged across horizon. We performed both a bivariate and partial correlation (correcting for age and IQ) using Pearson correlation.

### Computational modeling

We compared a set of generative models which assumed different exploration strategies accounting for subjects’ behaviour. Three core models were examined: UCB, Thompson sampling and a hybrid of those two. The UCB model captures directed and random exploration (the latter captured by a free softmax decision temperature parameter). The Thompson model reflects an uncertainty-dependent exploration, essentially a more complex softmax decision function. The hybrid model combines all of the above. We computed three extensions of each model by either adding tabula-rasa exploration, novelty exploration or both heuristics, leading to a total of 12 models (cf. Supplementary Material for details about the models).

## Results

### Subjects increase exploration when information can subsequently be exploited

To induce different extents of exploration, we manipulated the number of apples to be picked in each trial (termed ‘horizon’, one in the short horizon, six in the long horizon; Dubois et al. 2020). This horizon-manipulation promotes exploration in the long horizon because gaining new information can improve later choices.

To assess whether a longer decision horizon promoted exploration in our task, we compared which bandit subjects chose in their first draw in the short and in the long horizon condition. For each trial we computed the familiarity (the mean number of initial samples shown) and the expected value (the mean value of initial samples shown) of each bandit. In the long horizon condition, subjects preferred less familiar bandits (horizon main effect: F(1, 94)=5.824, p=.018, η^2^=.058; age main effect: F(2, 94)=.306, p=.737, η^2^=.006; age-by-horizon interaction: F(2, 94)=.836, p=.436, η^2^=.017; Fig. 2a), even at the expense of it having a lower expected value (horizon main effect: F(1, 94)=11.857, p=.001, η^2^=.112; age main effect: F(2, 94)=2.389, p=.097, η^2^=.048; age-by-horizon interaction: F(2, 94)=.031, p=.969, η^2^=.001; Fig. 2b). This is mainly driven by the fact that subjects selected the high-value bandit (i.e. the bandit with the highest expected reward based on the initial samples) less often in the long horizon (horizon main effect: F(1, 94)=24.315, p<.001, η^2^=.206; age main effect: F(2, 94)=1.627, p=.202, η^2^=.033; age-by-horizon interaction: F(2, 94)=2.413, p=.095, η^2^=.049; Fig. 4a), demonstrating a reduction in exploitation when information is useful. This behaviour resulted in a lower initial reward (on the 1^st^ sample) in the long compared to the short horizon (1^st^ sample: horizon main effect: F(1, 94)=13.874, p<.001, η^2^=.129; age main effect: F(2, 94)=1.752, p=.179, η^2^=.036; age-by-horizon interaction: F(2, 94)=1.167, p=.316, η^2^=.024; Fig. 2c).

**Fig. 2.**
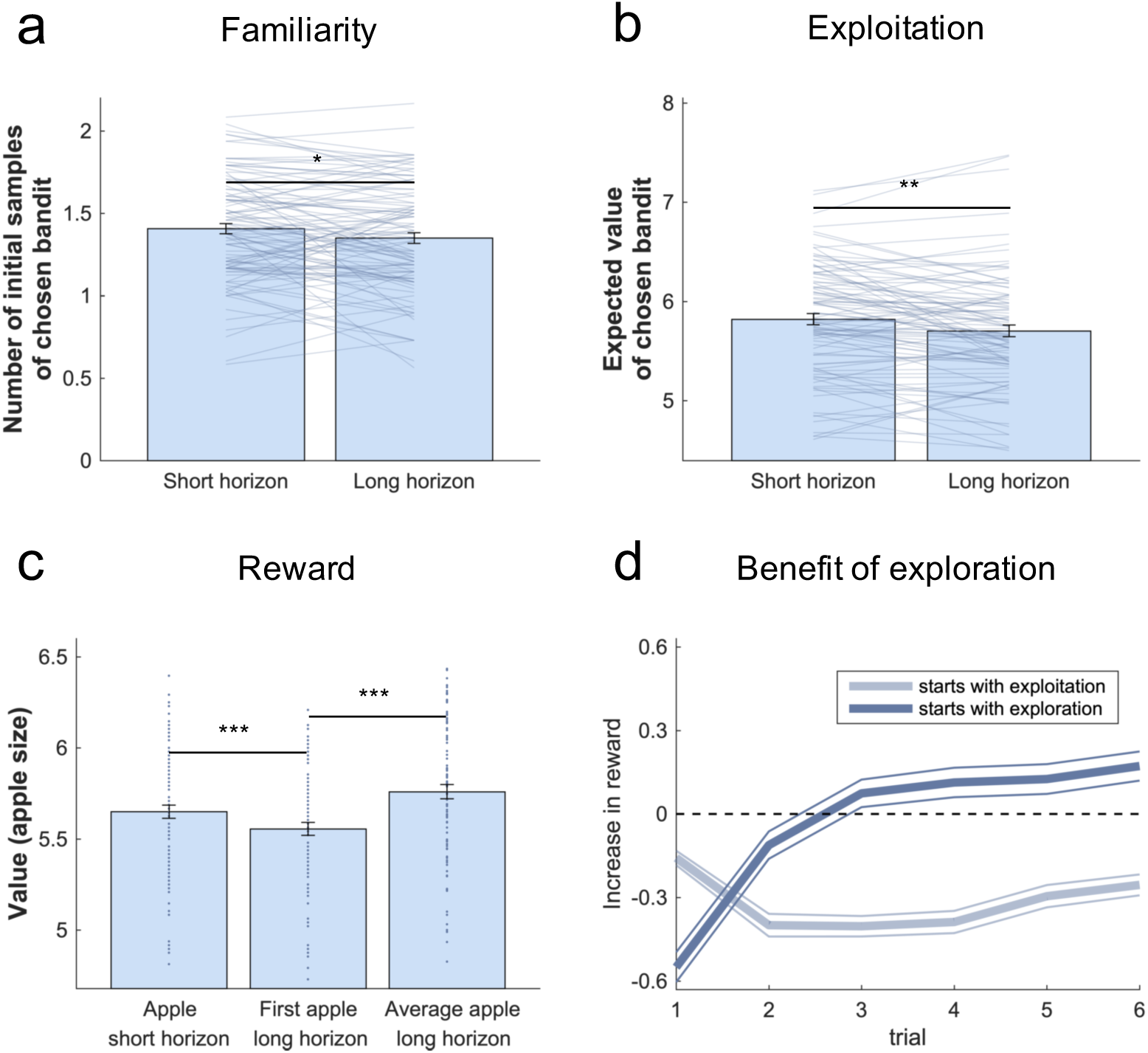
Benefits of exploration. (a) Subjects (collapsed across all age groups) chose less familiar (i.e. more informative) bandits more often on their first choice in the long compared to the short horizon. (b) Subjects chose bandits with a lower expected value (i.e. they exploited less) in the long horizon compared to the short horizon. (c) This behaviour led to a lower reward in the long horizon than in the short horizon on their first draw, indicating that subjects sacrificed larger initial outcomes for the benefit of more information. This additional information helped making better decisions in the long run, leading to higher earnings over all draws in the long horizon (right bar plot). Similarly, (d) in the long horizon, starting off by exploring (dark blue) versus exploiting (choosing the bandit with the highest expected value; light blue), led to an initial decrease in reward (negative increase in reward; difference between obtained reward and highest reward of initial samples), but eventually increased it. This means that the information that was gained through exploration led to higher long-term outcomes. * =p<.05, ** =p<.01, *** =p<.001. Data is shown as mean ± SEM and each dot/line represents a subject.

To evaluate whether subjects used the additional information in the long horizon condition beneficially, we compared the average reward (across six draws) obtained in the long compared to short horizon (one draw). The average reward was higher in the long horizon (horizon main effect: F(1, 94)=17.757, p<.001, η^2^=.159; age main effect: F(2, 94)=2.945, p=.057, η^2^=.059; age-by-horizon interaction: F(2, 94)=.555, p=.576, η^2^=.012; Fig. 2c), indicating subjects tended to choose less optimal bandits at first but subsequently made use of the harvested information to guide a choice of better bandits in the long run. This was also the case when we looked at the long horizon exclusively and compared the increase in reward (difference between the obtained reward and the highest shown reward) between when subjects started with an exploitative choice (chose the bandit with the highest expected value) versus an exploratory one. Exploration decreased their reward at first (exploration main effect: F(1, 94)=39.386, p<.001, η^2^=.295; age main effect: F(2, 94)=.443, p=.643, η^2^=.009; age-by-exploration interaction: F(2, 94)=.433, p=.650, η^2^=.009; Fig. 2d), but eventually increased it (6^th^ trial: exploration main effect: F(1, 94)=63.830, p<.001, η^2^=.404; age main effect: F(2, 94)=1.820, p=.168, η^2^=.037; age-by-exploration interaction: F(2, 94)=.753, p=.474, η^2^=.016; Fig. 2d), indicating that they were able to take advantage of the information gained through exploration.

### Subjects explore using computationally expensive strategies and simple heuristics

To determine which exploration strategies subjects use, we compared 12 models (cf. Supplementary Materials) using cross-validation, an approach that adequately arbitrates between model accuracy and complexity by comparing the likelihood of held-out data across different models. We compared a UCB model (directed exploration and random exploration), a Thompson model (uncertainty-dependent exploration), a hybrid of both and a combination of those with an *ϵ*-greedy (tabula-rasa exploration) and/or a novelty bonus (novelty exploration). These models made different predictions about how an agent explores and makes the first draw in each trial. Using Thompson sampling (Gershman 2018; Thompson 1933; captured by the Thompson model), she takes both expected value and uncertainty into account, with higher uncertainty leading to more exploration (uncertainty-dependent exploration). Using the UCB algorithm (Auer 2003; Gershman 2018; part of the UCB model), she also takes both into account but chooses the bandit with the highest (additive) combination of expected information gain and reward value (directed exploration). This computation is then passed through a softmax decision function (random exploration). The novelty bonus is a simplified version of the information bonus in UCB, which only applies to entirely novel options (novelty exploration). Using *ϵ*-greedy, a bandit is chosen entirely randomly, irrespective of expected values and uncertainties (tabula-rasa exploration). Just as in adults (Dubois et al. 2020), the winning model (Fig. 3a; cf. Supplementary Materials for more detail) in all three age groups combines Thompson sampling, *ϵ*-greedy and the novelty bonus, essentially a mixture of a complex (uncertainty-dependent) exploration, tabula-rasa exploration and novelty exploration. Simulations revealed that the winning model’s parameter estimates could be accurately recovered (Fig. 3b).

**Fig. 3.**
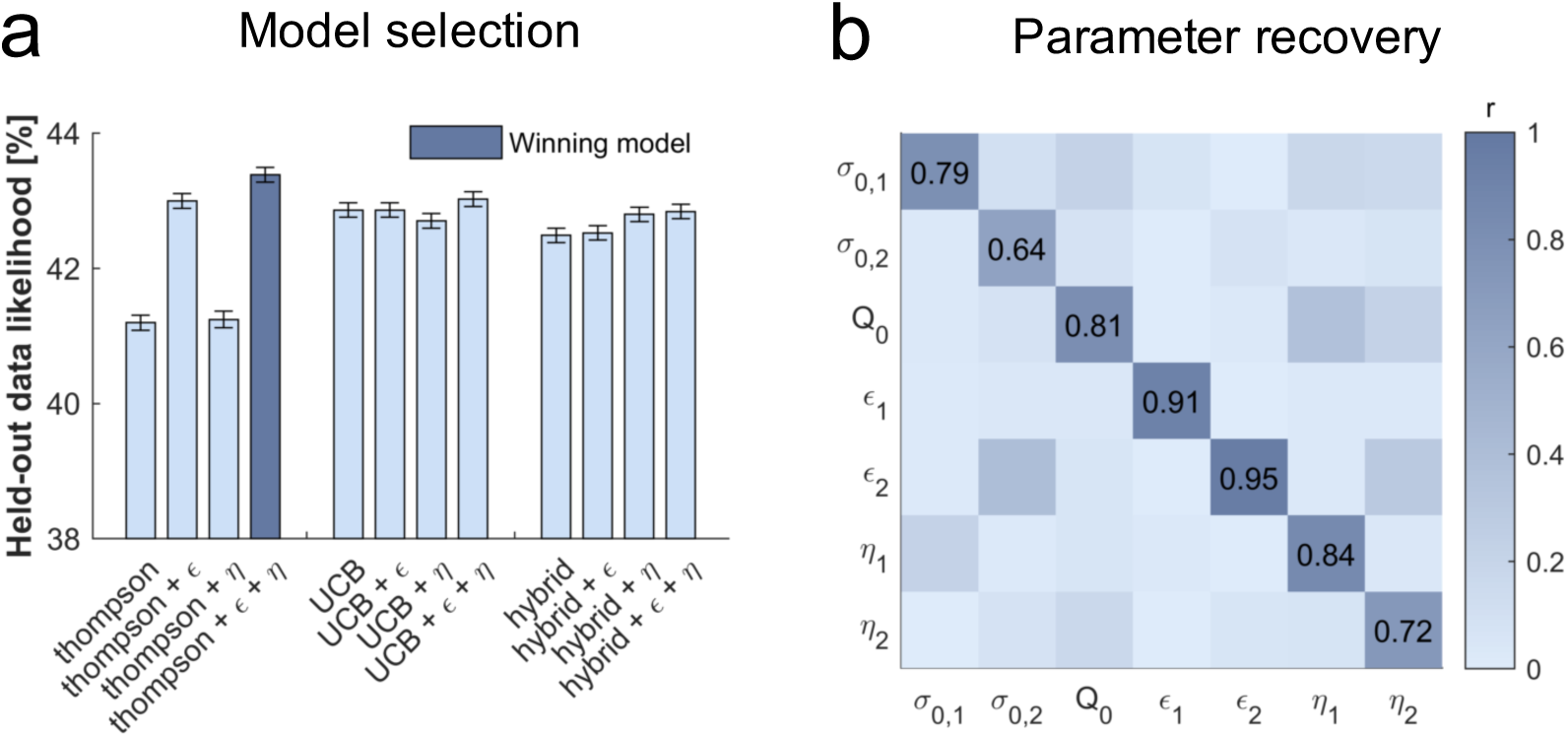
Subjects use a mixture of exploration strategies. (a) A 6-fold cross-validation of the likelihood of held-out data was used for model selection (chance level =33.3%). The Thompson model with both the tabula-rasa *ϵ*-greedy parameter and the novelty bonus *η* best predicted held-out data. (b) Model simulation with 4^7^ simulations predicted good recoverability of model parameters; *σ*_0_ is the prior variance and *Q*_0_ is the prior mean (cf. Supplementary Material for details about the models); 1 stands for short horizon-, and 2 for long horizon-specific parameters.

### Tabula-rasa exploration decreases in old adolescents

Tabula-rasa exploration (captured by *ϵ* -greedy) predicts that *ϵ* % of the time each option will have equal probability of being chosen. Under this regime, in contrast to other exploration strategies, bandits with a known low value are more likely to be chosen. This behavioural signature, the frequency of selecting the low-value bandit, was higher in the long compared to the short horizon condition (horizon main effect: F(1, 94)=8.837, p=.004, η^2^=.086; Fig. 4b). This was also captured more formally by analysing the *ϵ* parameter fitted values, which were larger in the long compared to the short horizon (horizon main effect: F(1, 94)=20.63, p<.001, η^2^=.180; Fig. 5a). These results indicate that subject made use of tabula-rasa exploration in a goal-directed way.

**Fig. 4.**
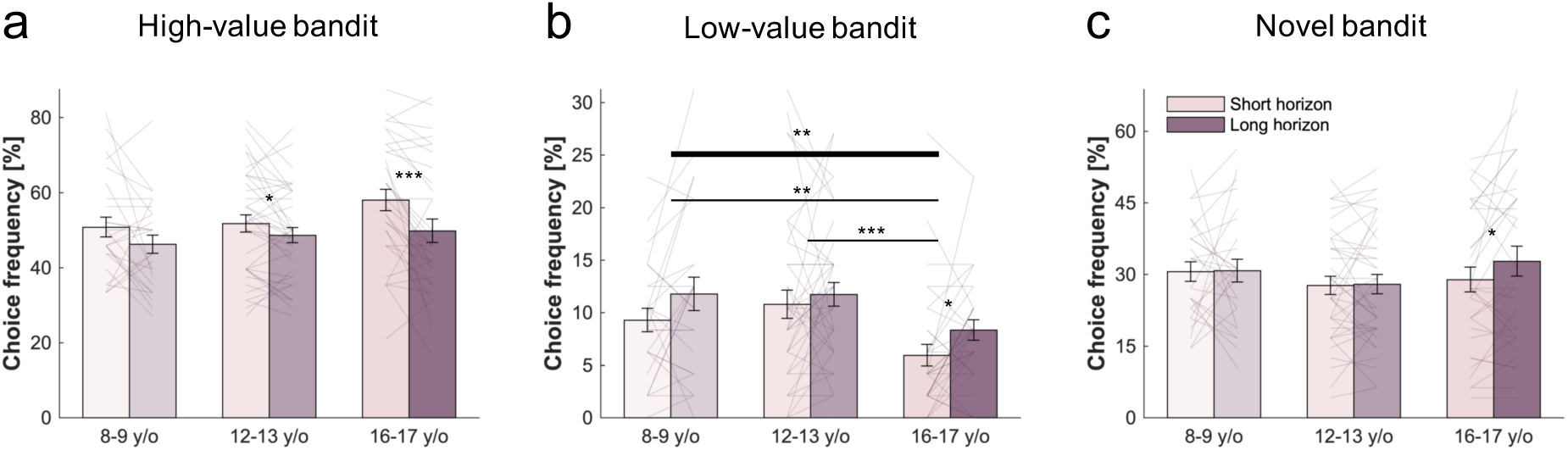
Behavioural age effects. Choice patterns in the first draw for each horizon and age group (children: 8-9 y/o, young adolescents: 12-13 y/o, old adolescents: 16-17 y/o). (a) Young and old adolescents, but not children, sampled from the high-value bandit (i.e. bandit with the highest average reward of initial samples) more in the short horizon compared to the long horizon, showing that the horizon manipulation altered exploration behaviour. (b) Old adolescents sampled less from the low-valued bandit compared to the children and young adolescents indicating that tabula-rasa exploration is reduced midway through adolescence. (c) Age groups did not differ in the amount of novelty exploration as measured by the choice frequency of the novel bandit, although it seems that its modulation by the horizon emerges in the old adolescent group. Horizontal bars represent rm-ANOVA (thick) and pairwise comparisons (thin). * =p<.05, ** =p<.01, *** =p<.001. Data is shown as mean ± SEM and each line represents one subject.

**Fig. 5.**
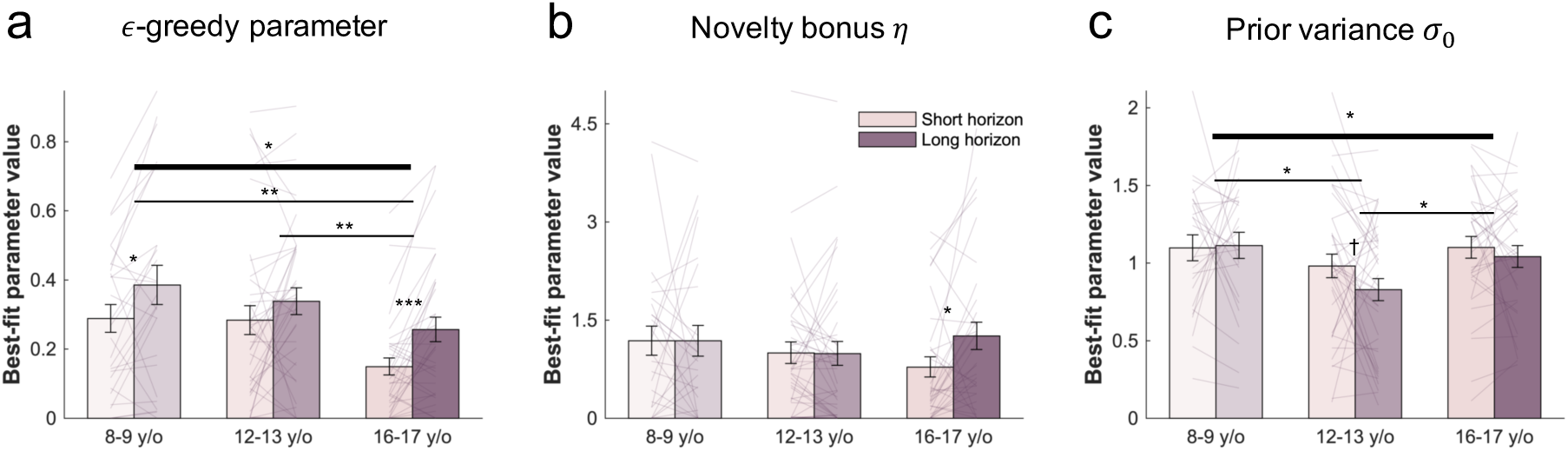
Age effects on model parameters. The winning model’s parameters were fitted to each subject’s first draw. (a) Old adolescents had lower values of *ϵ* (tabula-rasa exploration) overall compared to children and young adolescents, indicating that tabula-rasa exploration decreases during adolescence. Children and old adolescents had higher values of *ϵ* in the long compared to the short horizon. Subjects from all groups (b) assigned a similar value to novelty, captured by the novelty bonus η. It was higher (more novelty exploration) in the long compared to the short horizon for old adolescents indicating a goal-directed novelty exploration. (c) Young adolescents were less uncertain (lower prior variance *σ*_0_) about a bandits’ prior mean compared to children and old adolescents indicating a higher confidence and suggests a lower uncertainty-dependent exploration. † =p<.10, * =p<.05, ** =p<.01, *** =p<.001. Data is shown as mean ± SEM and each line represents one subject.

Next, we investigated whether the different age groups differed in their use of exploration strategies. We observed an age-related change in the use of tabula-rasa exploration. We found that age groups differed in the frequency of selecting the low-value bandit (age main effect: F(2, 94)=4.927, p=.009, η^2^=.095; age-by-horizon interaction: F(2, 94)=.236, p=.790, η^2^=.005; Fig. 4b). Interestingly, we found that the effect was primarily driven by a reduced tabula rasa exploration by old adolescents, compared to young adolescents and children (children vs old adolescents: t(52)=2.842, p=.005, d=.54; young vs old adolescents: t(76)=3.842, p<.001, d=.634), whilst children and young adolescents did not differ (t(52)=-.648, p=.518, d=.115). This suggests that the reduction in tabula-rasa heuristic usage occurs only later in adolescent development.

The same effect was observed when analysing the *ϵ* model parameter (age main effect: F(2, 94)=3.702, p=.028, η^2^=.073; age-by-horizon interaction: F(2, 94)=.807, p=.449, η^2^=.017; Fig. 5a). Again, this was driven by a reduced *ϵ* in the old adolescents compared to the younger groups (t(52)=3.229, p=.002, d=.622; young vs old adolescents: t(76)=2.982, p=.003, d=.491; children vs young adolescents: t(52)=.581, p=.562, d=.105). Our findings thus suggest that, compared to old adolescents, children and young adolescents rely more strongly on the computationally simple tabula-rasa exploration.

### No observed age effect on novelty exploration

We did not observe any difference for the novelty exploration strategy. There was no difference in the frequency of selecting the novel bandit (age main effect: F(2, 94)=.341, p=.712, η^2^=.007; horizon main effect: F(1, 94)=1.534, p=.219, η^2^=.016; age-by-horizon interaction: F(2, 94)=1.522, p=.224, η^2^=.031; Fig. 4c), nor more formally in the fitted novelty bonus *η* (age main effect: F(2, 94)=.341, p=.712, η^2^=.007; age-by-horizon interaction: F(2, 94)=2.119, p=.126, η^2^=.043; horizon main effect: F(1, 94)=1.892, p=.172, η^2^=.020; Fig. 5b). Interestingly, although not reaching statistical significance, it seems that unlike in children and young adolescents, old adolescents modulated their novelty exploration by the horizon. This can be observed both in the model parameter (*η*: horizon effect: old adolescents: t(33)=2.516, p=.017, d=.438; children: t(26)=-.003, p=.998, d=.001; young adolescents: t(38)=-.079, p=.937, d=.013; Fig. 5b) and in behaviour (frequency of selecting novel bandit: horizon effect: old adolescents: t(33)=2.199, p=.035, d=.383; children: t(26)<.001, p=1, d<.001; young adolescents: t(38)=.04, p=.969, d=.006; Fig. 4c), indicating that subjects seem to become more goal-directed during development.

### Young adolescents use less uncertainty-dependent exploration

In addition to heuristics, we found that all age groups used complex exploration strategies to some extent. Those strategies rely on the computation of expected values and uncertainties. To assess whether different age groups used those complex strategies differently, we compared the model-derived prior mean *Q*_0_ and prior variance (or uncertainty) *σ*_0_, which are used respectively for the computation of the expected value and the uncertainty of each bandit (for details about the models cf. Dubois et al. 2020; Gershman 2018). Essentially, *Q*_0_ is the reward that subjects expect to get from a bandit before integrating its initial samples, and *σ*_0_ is their uncertainty about this *Q*_0_. The prior mean *Q*_0_ was similar in all groups (F(2,94)=1.367, p=.260, η^2^ =.028). However, a difference in the prior variance *σ*_0_ (i.e. uncertainty) was observed (age main effect: F(2, 94)=3.241, p=.044, η^2^=.065; age-by-horizon interaction: F(2, 94)=.866, p=.424, η^2^=.018; horizon main effect: F(1, 94)=1.576, p=.212, η^2^=.016; Fig. 5c) indicating differences in uncertainty at the beginning of a trial. This was driven by the young adolescents having a lower *σ*_0_ (less uncertainty) compared to children and old adolescents (pairwise comparisons: children vs young adolescents: t(52)=2.55, p=.012, d=.452; young vs old adolescents: t(76)=-2.309, p=.022, d=.38; children vs old adolescents: t(52)=.445, p=.657, d=.0835). Taken together, those results indicate that before integrating the initial samples, young adolescent were more certain in their belief about the reward value of future samples, suggesting less uncertainty-dependent exploration.

### Tabula-rasa exploration is linked to ADHD symptoms

Developmental effects on exploration strategies are important to understand developmental psychiatric disorders, such as ADHD, which has been suggested to be linked to excessive exploratory behaviour (Hauser et al. 2014, 2016). We have previously shown that tabula-rasa exploration is modulated by noradrenaline (Dubois et al. 2020). As this neurotransmitter is thought to be disrupted in ADHD (Arnsten and Pliszka 2011; Berridge and Devilbiss 2011; Del Campo et al. 2011; Frank et al. 2007; Hauser et al. 2016; Luman, Tripp, and Scheres 2010), we hypothesized that tabula-rasa exploration would be linked to ADHD symptoms. We thus compared whether the amount of tabula-rasa exploration was associated with ADHD scores. We found that ADHD symptoms were significantly associated with tabula-rasa exploration captured by the model parameter *ϵ* (*r =*.260, p=.010; Fig. 6) and as indicated by the low-value bandit picking frequency (*r =*. 259, p=.010). The effect remained when additionally controlling for age and IQ (partial correlation with *ϵ*: *r*=.218, p=.034; with low-value bandit picking: *r*=.214, p=.037). ADHD symptom did not correlate with any of the other exploration strategies (with *η*: *r* = −.113, p=.269; with *σ*_0_: *r* = −.005, p=.963), suggesting that tabula-rasa is the most relevant exploration factor for ADHD symptoms.

**Fig. 6.**
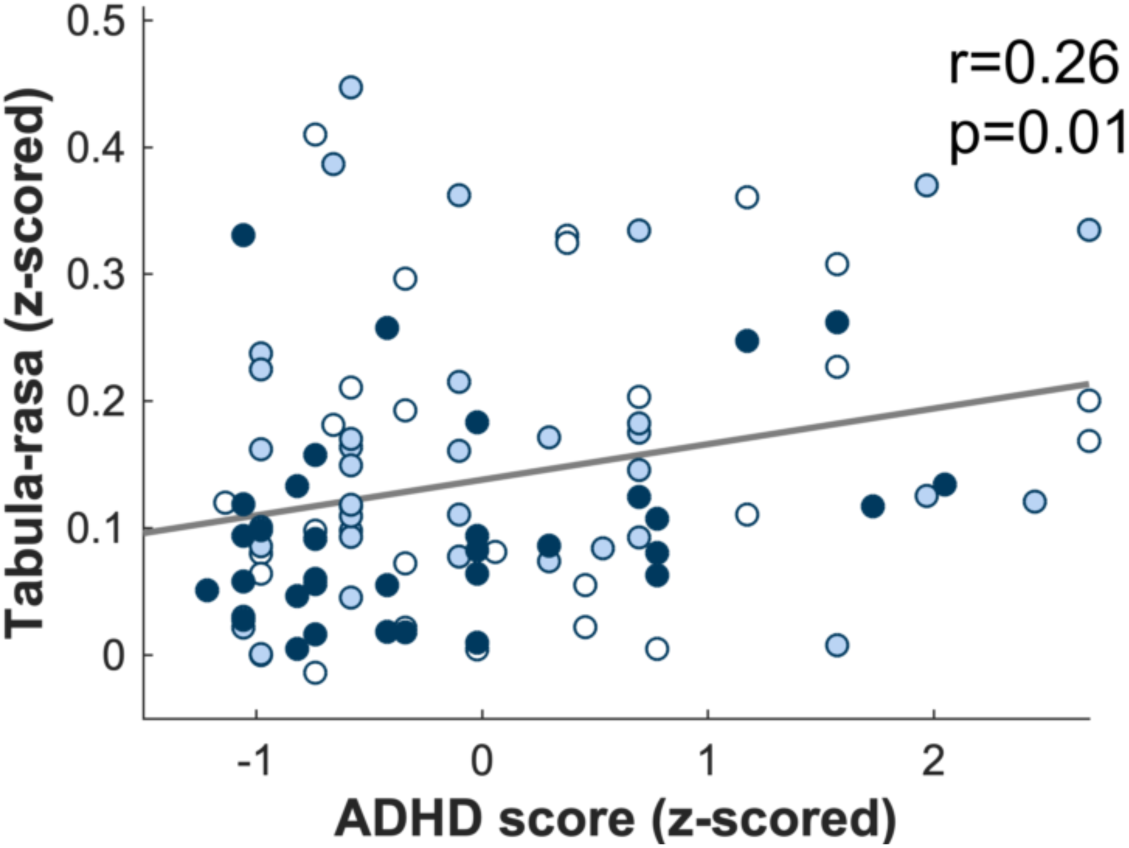
Tabula-rasa exploration increased in subjects with high ADHD scores. Tabula-rasa exploration (as captured by the model parameter *ϵ*) correlated with ADHD scores indicating that it is specifically this exploration strategy which is impaired in ADHD. Each dot represents one subject. White: children, light blue: young adolescents, dark blue: old adolescents.

## Discussion

Given limited cognitive and neural resources, how is it possible that young people are able to solve the complex and computationally demanding exploration-exploitation trade-off so successfully, and to learn at an unprecedented pace? We have previously shown that even adults depend on simple exploration heuristics to bypass expensive computations (Dubois et al. 2020). In this study we show that children and young adolescents rely more heavily on these exploration heuristics, in particular tabula-rasa exploration, when balancing between exploiting known and exploring less well-known options.

By assigning the same choice probability to all options, effectively suppressing the need to keep track of any expected values, tabula-rasa exploration requires minimal computational resources. However, this computational efficiency comes as the cost of choice sub-optimality, due to the occasional selection of options of low expected value. Despite its sub-optimality, we demonstrate that this heuristic is more intensely used at an early age. Our findings thus suggest that the reduced cognitive resources during childhood are likely to be accommodated using computationally less demanding exploration strategies. Moreover, younger individuals may not be that negatively affected by the limitations of tabula-rasa exploration because their limited (life) experience has not yet allowed them to build sophisticated models of the world. Specifically, the limited learning experiences means that their beliefs are more imprecise or even inaccurate. Therefore, ignoring those weak and unstable priors does not significantly penalize learning. It may even help prevent the integration of (initially) falsely rated states, essentially accounting for childrens’ erroneous believes due to lack of experience.

Interestingly, it seems that the transition in exploration strategies is not a continuous process during development but occurs mid-way through adolescence. Children (8-9 y/o) and young adolescents (12-13 y/o) show similar patterns of exploration, whereas the old adolescent (16-17 y/o) group differed from both younger ones. The two younger groups employ more tabula-rasa exploration compared to the older group. Additionally, novelty exploration seems to become horizon-dependent in the older group, similarly to what is observed in adults (Dubois et al. 2020), suggesting an emergence of a goal-directed novelty exploration around this period. This adds to the number of cognitive abilities that improve during adolescence (Gopnik 2020; Luna et al. 2004; Waber et al. 2007) and corresponds to brain maturation around that period (Geidd 2004; Giorgio et al. 2010; Gogtay et al. 2004; Tamnes et al. 2010), in particular the PFC (Casey et al. 2005; Segalowitz and Davies 2004), which is essential in integrating complex sources of information required for advanced decision-making (Hartley and Somerville 2015). This late maturation, corresponding to an increase in complex information integration (Chrysikou et al. 2013; Gopnik, Griffiths, and Lucas 2015), is thought to be responsible for the slow calibration of executive function during development (Anderson 2002; Blakemore and Choudhury 2006; Diamond 2009). As those regions are less accessible or not functioning optimally in younger individuals, they might circumvent this problem by the use of less-resource demanding strategies (i.e. heuristics) for exploration, and switch to more complex strategies as they grow older.

In addition to heuristics usage, development seems to impact confidence in prior knowledge. Young adolescents were more certain about their initial beliefs compared to both children and old adolescents. This is line with studies demonstrating a hyper-confidence in young adolescents (Moses-Payne et al. *submitted*), and may imply a lower curiosity (Wade and Kidd 2019). Additionally, as uncertainty increases exploration when complex strategies (e.g. Thompson sampling and UCB) are used, our findings suggest less resource-demanding exploration at the beginning of adolescence.

We found that tabula-rasa exploration was more present in subjects with increased ADHD symptoms, irrespective of their age. ADHD is believed to be linked to an impairment in the dopaminergic and noradrenergic systems (Arnsten and Pliszka 2011; Berridge and Devilbiss 2011; Del Campo et al. 2011; Frank et al. 2007; Hauser et al. 2016; Luman, Tripp, and Scheres 2010) with common ADHD medication targeting dopamine (e.g. methylphenidate; Iversen 2006) and noradrenaline functioning (e.g. atomoxetine; Levy 2008). We have previously demonstrated that tabula-rasa exploration is modulated by noradrenaline (Dubois et al. 2020), and interestingly it is specifically this form of exploration which is associated with ADHD in our study. This suggests that it might be the impairment in noradrenaline which underlies the increase of tabula-rasa exploration in ADHD. Our findings thus also extend previous work (Hauser et al. 2014), where an altered exploration-related behaviour was found in adolescents with ADHD, but it was unclear which type of exploration or which type of neurotransmitter was affected. This finding helps understand ADHD symptomatology, both from a computational and evolutionary perspective, i.e. in what environment it can be adaptive (Williams and Taylor 2006). Future studies could make use of a similar paradigm in a longitudinal setup in order to understand the individual trajectories of exploration and how ADHD symptoms and tabula-rasa exploration influence each other.

Exploration is an essential part of learning (Gopnik 2020; Kidd and Hayden 2015; Sutton and Barto 1998) and thus crucial for development. In this study we show substantial changes in exploration between childhood and adolescence. Our results thus clearly expand the previous studies that either only focused on exploration in children (Bonawitz et al. 2011, 2012; Cook, Goodman, and Schulz 2011; Gweon et al. 2014; Meder et al. 2020; Pelz, Yung, and Kidd 2015), or compared adults to minors without separating development within childhood and adolescence (Schulz et al. 2019; Somerville et al. 2017). Moreover, to our knowledge we are the first to compare between multiple computationally complex and simple exploration strategies and show a specific development of tabula-rasa exploration in the transition from childhood to adolescence. Our findings thus demonstrate that youths deploy a multitude of exploration strategies, and that the reliance on these strategies dynamically changes before reaching adulthood.

Taken together our results suggest that tabula-rasa exploration is a simple exploration strategy that is of great benefit when cognitive resources are still limited during earlier stages of development. As we grow older, and our experience expands, it is evolutionary useful to incorporate our knowledge in decision making, and therefore lessen their use of tabula-rasa exploration. Such a process may go awry in people with ADHD symptoms, as they show increased levels of tabula-rasa exploration, which could, in the long-term, lead to suboptimal decision making.

## Conflict of interest

The authors declare no competing financial interests.

## Acknowledgements

The authors thank the children, adolescents, their schools (especially Sydney Russell School, Baden Powell School and Holmleigh Primary School) and families for taking part in this study. Thanks are also given to S. Waters for helping with data collection. M.D. is a predoctoral fellow of the International Max Planck Research School on Computational Methods in Psychiatry and Ageing Research. The participating institutions are the Max Planck Institute for Human Development and the University College London (UCL). T.U.H. is supported by a Wellcome Sir Henry Dale Fellowship (211155/Z/18/Z), a grant from the Jacobs Foundation (2017-1261-04), the Medical Research Foundation, and a 2018 NARSAD Young Investigator Grant (27023) from the Brain and Behavior Research Foundation. The Max Planck UCL Centre is a joint initiative supported by UCL and the Max Planck Society. The Wellcome Centre for Human Neuroimaging is supported by core funding from the Wellcome Trust (203147/Z/16/Z).

## Supplementary Material

### Computational modelling

The models used have been developed and discussed in detail in Dubois et al. 2020. Here we re-print these equations for completion.

The value of each bandit is represented as a distribution *N*(*Q, S*) with *S* = 0.8. Subjects have prior beliefs about bandits’ values which we assume to be Gaussian with mean *Q*_0_ (prior mean; free parameter) and uncertainty *σ*_0_ (prior variance; free parameter).

#### Mean and variance update rules

The expected mean *Q* and precision 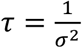 of each bandit are updated as follows:

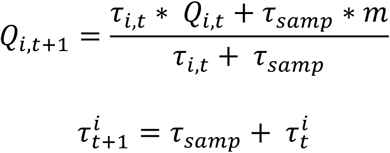

with 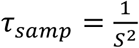 the sampling precision, *S* = 0.8 the fixed sampling variance, *m* the presented sample, *i* is the bandit and *t* the time point.

We examined three base models: UCB, Thompson sampling and a hybrid of UCB and Thompson sampling. We computed three extensions of each model by either adding tabula rasa exploration (c_tr_ c_n_) = {1,0}, novelty exploration (c_tr_ c_n_) = {0,1} or both heuristics (c_tr_ c_n_) = {1,1}, leading to a total of 12 models. For each model, the probability of choosing a bandit is described below.

#### UCB

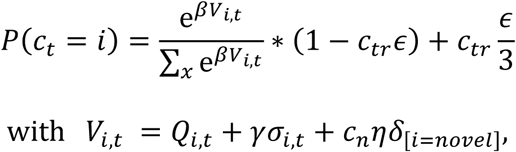

*γ* the information bonus, *V* the expected value, *η* the novelty bonus and *β* the inverse temperature.

#### Thompson sampling

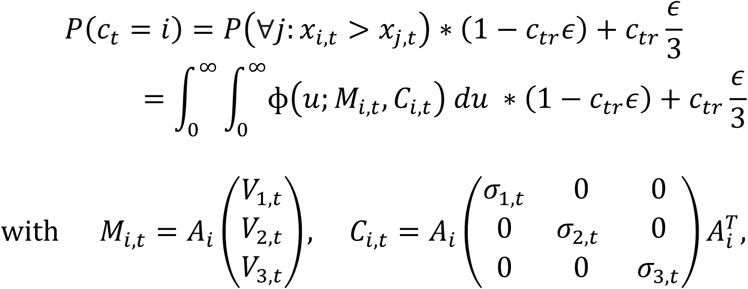

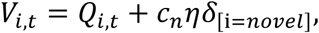

Φ the multivariate Normal density function, *A* the matrix computing the pairwise differences for each bandit and 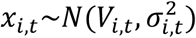 a sample taken from each bandit.

#### Hybrid model

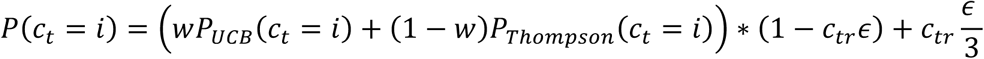

with *w* the contribution of each of the UCB and Thompson sampling models

#### Parameter estimation

To fit the parameter values, we used the maximum a posteriori probability (MAP) estimate. The optimisation function used was fmincon in MATLAB. The parameters could vary within the following bounds: *σ*_0_ = [10^−8^, 8], *Q*_0_ = [1, 10], *ϵ* = [0, 1], *η* = [0, 5]. The prior distribution used for the prior mean *Q*_0_ and the prior variance *σ*_0_ parameters were the normal distributions that approximate the generative distributions: *Q*_0_ ∼ *N*(5, 2) and *σ*_0_ ∼ *N*(1.4, 1). For the *ϵ*-greedy parameter and the novelty bonus *η* a uniform distribution was used (equivalent to performing maximum likelihood estimation).

#### Further modeling

Model comparison was performed using K-fold cross-validation (K=6). For details about model comparison, parameter recovery, model validation and free parameters, cf. Dubois et al. 2020.

**Table S1.** Characteristics of age groups. The age groups did not differ in gender, intellectual abilities (adapted WASI matrix test, adjusted for age) nor ADHD symptoms (Conners 3AI-SR, Conners 2008). Mean (SD).

|  | Children | Young adolescents | Old adolescents |  |
| --- | --- | --- | --- | --- |
| Age | 9.32 (.27) | 13.13 (.30) | 17.18 (.30) |  |
| Gender (M/F) | 10/16 | 17/21 | 14/19 | $F(2, 94)=.121, p=.886, \eta^2=.003$ |
| Intellectual abilities | 93.88 (12.88) | 98.50 (13.86) | 97.45 (10.47) | $F(2, 94)=1.095, p=.339, \eta^2=.023$ |
| ADHD symptoms | 59.23 (13.41) | 56.05 (12.90) | 54.18 (11.30) | $F(2, 94)=1.191, p=.308, \eta^2=.025$ |

## Notes

### Competing Interest Statement

The authors have declared no competing interest.

